# Metabolomic profiling of pancreatic adenocarcinoma reveals fundamental clinical features

**DOI:** 10.1101/2021.02.01.429087

**Authors:** Abdessamad El Kaoutari, Nicolas A Fraunhoffer, Owen Hoare, Carlos Teyssedou, Philippe Soubeyran, Odile Gayet, Julie Roques, Gwen Lomberk, Raul Urrutia, Nelson Dusetti, Juan Iovanna

## Abstract

In this study, we perform the metabolomics profiling of 77 PDAC patient-derived tumor xenografts (PDTX) to investigate the relationship of metabolic profiles with overall survival (OS) in PDAC patients, tumor phenotypes and resistance to five anticancer drugs (gemcitabine, oxaliplatin, docetaxel, SN-38 and 5-Fluorouracil). We identified a metabolic signature that was able to predict the clinical outcome of PDAC patients (p < 0.001, HR=2.68 [95% CI: 1.5-4.9]). The correlation analysis showed that this metabolomic signature was significantly correlated with the PDAC molecular gradient (PAMG) (R = 0.44 and p < 0.001) indicating significant association to the transcriptomic phenotypes of tumors. Resistance score established, based on growth rate inhibition metrics using 35 PDTX-derived primary cells, allowed to identify several metabolites related to drug resistance which was globally accompanied by accumulation of several diacy-phospholipids and decrease in lysophospholipids. Interestingly, targeting glycerophospholipid synthesis improved sensitivity to the three tested cytotoxic drugs indicating that interfering with metabolism could be a promising therapeutic strategy to overcome the challenging resistance of PDAC.

**Short abstract:** Targeting metabolism of cancer cells gives a precious opportunity to overcome challenges related to the high mortality and chemoresistance in PDAC.

Metabolic profiling of PDAC patient-derived tumor xenografts used in this study allowed highlighting the strong link between metabolism and both clinical outcome of the patients and chemoresistance.

Metabolic signature was able to discriminate between good and bad prognosis groups of patients based on their level of key metabolites.

Identification of key metabolic markers associated to chemoresistance allowed to improve sensitivity to anticancer drugs.

These results provide new insights to help to predict patient survival and elaborate new combinatory therapies against chemoresistance in PDAC patients attesting of the important clinical value of this work.

## Introduction

Pancreatic ductal adenocarcinoma (PDAC) is one of the most aggressive and lethal cancers with a dismal outcome due to many factors including the high heterogeneity of tumors, late diagnosis, and high resistance to chemotherapies. The overall 5-year survival rate is currently around 8% which highly depends on the surgery and stage of disease (Siegel et al, 2018). For example, a tumor resection combined with adjuvant therapies can increase this rate to 20% (Neoptolemos et al, 2017). However, only 15 to 20% of patients are potentially resectable at the time of diagnosis (Hartwig et al, 2013). PDAC tumors have been classified commonly into two main subtypes, classical, with a better prognosis, and basal-like with a poorer clinical outcome (Moffitt et al, 2015; Yachida & Iacobuzio-Donahue, 2013). However, the heterogeneity in PDAC tumors is higher than anticipated as shown in recent studies which indicate that not only basal-like and classical cell populations coexist in the same tumor (Juiz et al, 2020) but, a continuum distribution of phenotypes describing PDAC aggressiveness, rather than a binary system, is present in PDAC subtypes (Nicolle et al, 2020). The high level of heterogeneity in PDAC results from a combination of genetic, epigenetic, and micro-environmental alterations, which is directly related to development of drug resistance. In fact, during treatment, a small subpopulation of cancer cells may be able to metabolize the anticancer drug, thereby developing a resistance that allows them to grow and become the dominant population. This resistance to therapeutic molecules, either intrinsic or newly developed, is still the major challenge in PDAC treatment (Zeng et al, 2019). Indeed, investigations on the mechanisms underlying drug resistance over the past decade have contributed toward the better understanding of this disease, but more knowledge is needed to improve PDAC chemotherapy.

The metabolic activities in cancer cells are reprogrammed to meet the need of tumors for growth and rapid proliferation through increased biosynthesis and metabolism of macromolecules, such as lipids and amino acids (Cairns et al, 2011; Gunda et al, 2017; Tadros et al, 2017; Vander Heiden et al, 2009). Therefore, given that metabolism is a key regulator of tumorigenesis, therapeutic efficacy is related to the ability to modulate the metabolic alterations of tumor cells. As recently reported, metabolic alterations have been associated with drug resistance in cancer cells (Zong et al, 2018). In PDAC, the inhibition of fatty acid biosynthesis is able to overcome gemcitabine resistance (Tadros et al, 2017). Furthermore, other studies have shown that the oxidative phosphorylation (OXPHOS) pathway in mitochondria significantly contributes to drug resistance. In fact, chemotherapy resistance was associated to high OXPHOS status in several cancers (Bosc et al, 2017; Desbats et al, 2020). Targeting high OXPHOS in PDAC tumors is synergistic with gemcitabine treatment (Masoud et al, 2020), and disrupting lipid-rafts sensitizes resistant pancreatic tumor initiating cells to standard chemotherapy and decreases their metastatic potential (Gupta et al, 2018). Hence, therapeutic strategies based on the modulation of metabolism in combination with chemotherapeutic drugs provide a promising opportunity to overcome drug resistance in pancreatic cancer.

In this study, we investigated the metabolic profiles of 77 PDAC PDTXs (patient-derived tumor xenografts) and analyzed their relationship with the resistance to five anticancer drugs to identify relevant metabolic biomarkers. Notably, we found that some metabolites are associated to multidrug resistance, and through modifying the lipidomic profile, we were able to re-senstize PDAC cells to some cytotoxic drugs. Overall, our study reveals key therapeutic targets to improve the treatment of PDAC and overcome drug resistance.

## Results

### Metabolic profiling of PDAC

We performed metabolic profiling on 77 pancreatic cancer samples grown as PDTX. Unsupervised clustering analysis of these metabolic profiles revealed distinct clusters on the dendrogram, indicating key heterogeneity among the PDAC samples analyzed. The metabolic heterogeneity across samples was related to the intensity variations in metabolites that belong to different classes as shown in Figure 1A. Considering the entire metabolome dataset of all samples, a total of 502 metabolites were detected and included in further analysis. The vast majority of these metabolites were from the lipid class. Glycerophospholipids were the most represented class with 45% metabolites followed by glycerolipids (17.1%), fatty acids (10%), sphingolipids (8.2%), amino acids (5.8%), nucleotides (2.8%) and sterols (2.2%), as summarized in Figure 1B. The other metabolites, representing 9% of all metabolites, included carbohydrates, such as monosaccharides and disaccharides, as well as alkylamines. All metabolite data corresponding to each patient is shown Dataset S1 and the annotation of metabolic feature in Dataset S2. Thus, the metabolomic profiles of human PDAC tumors grown as PDTXs display heterogeneity that is comprised largely of the lipid class of metabolites.

**Figure 1.**
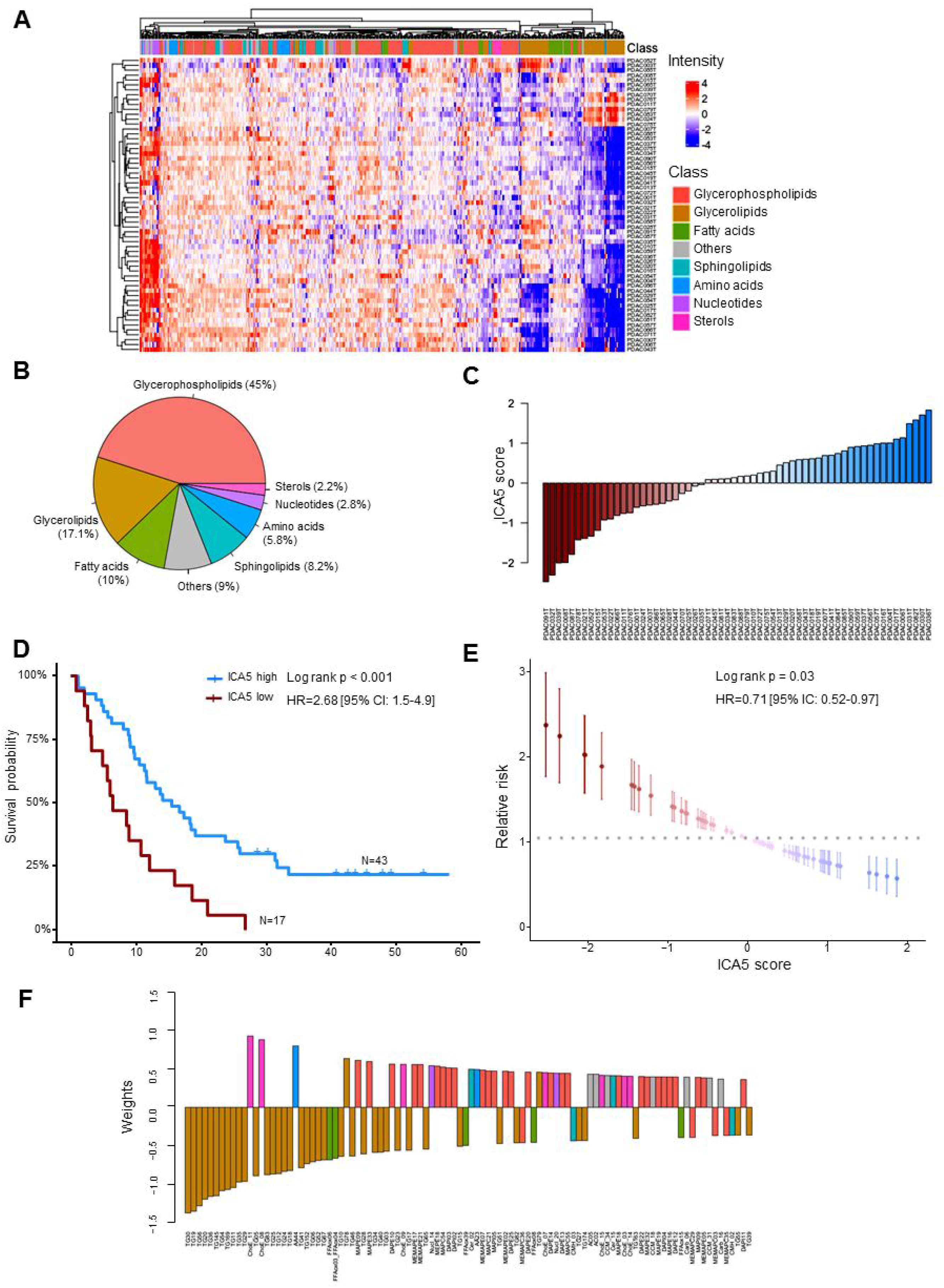
Metabolic profiling and prognostic signature of PDAC patients. A) Heatmap showing differential metabolic profiles of PDAC samples using unsupervised hierarchical clustering. The column annotation represents the main classes of metabolites. B) Pie chart indicating the global distribution of metabolic classes in all samples. C) Waterfall plot showing the sorted scores of patients in the identified metabolic signatures. D) Kaplan-Meier plot of survival using univariate analysis based on ICA5 scores of patients. Two groups of patients were considered with high and low score. E) Univariate relative risk for overall survival associated with the metabolic signature. Each point is a patient’s relative risk of disease with error bars corresponding to a 95% confidence interval. F) Barplot illustrating the contribution of the top 100 metabolites into the component ICA5.

### New metabolic signature to predict patient survival in PDAC

To investigate whether a metabolic profile could impact the survival of PDAC patients, we performed an independent component analysis including all the metabolites, followed by a survival analysis using univariate Cox regression based on PaCaOmics patient cohort (Nicolle et al, 2020) to assess their prognostic value. ICA5 component significantly associated with overall survival (OS) in our patient cohort. The weights in this component attributed a score for each sample in the dataset and subsequently used for survival analysis by creating low and high score groups (Figure 1C). Patient survival in the high-ICA5-score group was significantly improved compared with patients in the low-ICA5-score group with log rank p-value < 0.001 and hazard ratio HR = 2.68 (95% CI: 1.5-4.9) (Figure 1D). These results demonstrated that the metabolic signature identified here has a significant predictive value in OS of patients with PDAC. This predictive value of the metabolic component was then confirmed using a relative risk (RR) regression model. Results showed that patients with lower ICA5 score have significantly higher RR (log rank p = 0.03, HR = 0.71 [95% IC: 0.52-0.97]) than patients with higher ICA5 score (Figure 1E). Clinical features and the ICA5 score of each patient are presented in the Dataset S3.

As described above, the most represented metabolites in the overall patient metabolome belonged to the glycerophospholipid class; however glycerolipids contributed most to the metabolomics signature identified here for predicting patient survival. In Figure 1F, we represented the top 100 metabolites according to their weights (negative or positive contribution) in component ICA5. Triacylglycerols, such as TG30 and TG56 along with several oxidized fatty acids, had a negative association to the component suggesting that these metabolites could be associated with poor prognosis. However, cholesteryl esters (ChoE_11 and ChoE_8) and glycerophospholipids, especially lysophospholipids, were positively associated to the component indicating a possible link with improved prognosis in patients with high ICA5 scores (Figure 1F and Dataset S4). Therefore, component analysis of metabolomic profiles allows us to idenfity key signature associated with patient survival.

### The metabolomics profile associated with phenotype of PDAC

To investigate the link between the metabolomic profiles and molecular phenotypes, we compared the metabolic signature identified in this study with the PDAC molecular gradient (PAMG) previously described (Nicolle et al, 2020). The PAMG is a transcriptomic signature that describes PDAC heterogeneity as a continuous gradient from pure basal-like (low PAMG) to pure classical phenotypes (high PAMG). The correlation analysis showed that the metabolic signature identified in this study significantly correlated with the PAMG with a *Pearson* coefficient R = 0.44 and p < 0.001 (Figures 2A and 2B), indicating the association between the metabolic signatures and transcriptomic phenotypes of the tumors. Along with their good prognosis estimation, tumors with high ICA5 scores appeared to be more classical, while tumors with low ICA5 scores were characterized by a basal-like phenotype with poor prognosis. These results show that the metabolic signature identified here is strongly associated with both tumor phenotypes and OS of patients in PDAC. Furthermore, we performed correlation analysis to identify all metabolites that could be associated individually with the PAMG. Globally, out of 502 metabolic features analyzed, 97 positively correlated with the PAMG and were mainly glycerophospholipids (70%), while 88 metabolites negatively correlated and belonged mainly to glycerolipids (Figure 2C). For instance, lyso-phosphatidylcholines (LPC) and lyso-phosphatidylethanolamine (LPE), which are part of the glycerophospholipids class, correlated positively to the PAMG, whereas triacylglycerols (TAGs), which are classified as glycerolipids, such as TG73 and TG68, were inversely correlated with the PAMG. As much as 65% of the negatively-correlated metabolites were glycerolipids, followed by glycerophospholipids (15%), in particular 2-acyl-glycero phosphatidylcholine (MEMAPC), and sphingolipids (12.5%) as presented in Figure 2C). Overall, these results display the strong association between transcriptomic heterogeneity and the metabolic heterogeneity in PDAC patients. We conclude that the more basal-like a tumor is the higher likelihood it exhibits increased levels of fat lipids, such triacylglycerols, as opposed to a classical tumor that would be more prone to present with an accumulation of glycerophospholipids, such as LPC and LPE. The increased level of TAGs in basal-like tumors could be characteristic of tumor aggressiveness, as these fat lipids are the main components of lipid droplets that constitute an important reservoir of fatty acids and energy for cell growth and proliferation.

**Figure 2.**
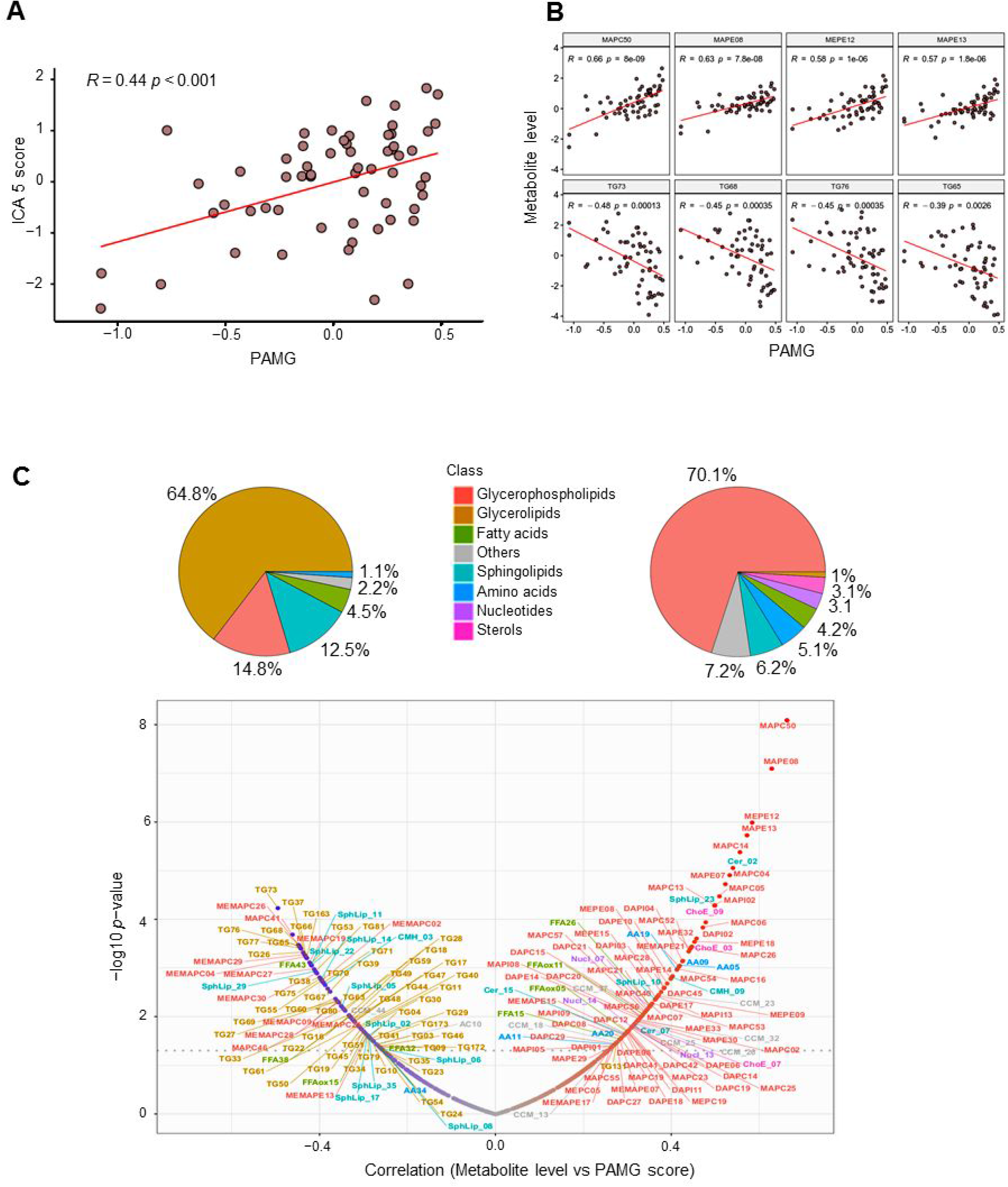
Metabolic signature is correlated with molecular gradient of the tumors. A) Scatter plot showing the correlation between molecular gradient and ICA5 scores (*Pearson* coeficient 0.38 and p = 0.0031). B) Scatter plots of most correlated metabolites to molecular gradient. Statistics of the *Pearson’s* correlation are shown. C) Visualization of all correlated metabolic features to molecular gradient. Pie charts and volcanoplot illustrate the repartition among main classes of anticorrelated (left) and correlated (right) metabolites. All metabolites with a *Pearson* correlation p values < 0.05 were shown.

### Establishment of drug resistance score

To explore the association between the metabolic profiles and drug resistance, we established a new score of resistance based on GR inhibition metrics as described in the methods section. In fact, traditional dose-response metrics such IC50 and Emax could be confounded by the number of cell divisions and the assay duration which induces bias to biological effect of the drug (Hafner et al, 2016). Using GR metrics fitted to sigmoidal curves allowed thereby a better quantification of the response to drugs. Dose-response data of 35 primary cell lines derived from pancreatic PDTX samples were used to calculate metrics for five drugs tested in this study including gemcitabine, oxaliplatin, docetaxel, SN-38 and 5-fluorouracil (5FU) (Figure 3A). For each drug, after performing principal component analysis (PCA), a weighting coefficient was calculated for each metric based on its correlation to the first three dimensions that explain more than 90% of variance in the dataset (Supplementary Figures S1 to S5). To standardize the score calculation, the average weighting coefficient across drugs was used to calculate the GRM score of resistance as follows: GRMs = (log10(GR50) x GR50Coeff) + (GRmax x GRmaxCoeff) + (GR_AOC x AOCcoeff) + (hGR x hCoeff). It is important to note that all used metrics have positive coefficients except AOC, which is negatively associated to other metrics and had negative weighting coefficient. Thus, GR50Coeff = 1.9, GRmaxCoeff = 2, AOCcoeff = -2.25 and hCoeff = 0.72. The GRM scores were then centered and scaled for further analysis. The score of each drug ranked the tumors from the most resistant (highest score) to the most sensitive by considering multiple metrics normalized to growth rate to provide more robust estimation of drug resistance (Figure 3B to F).

**Figure 3.**
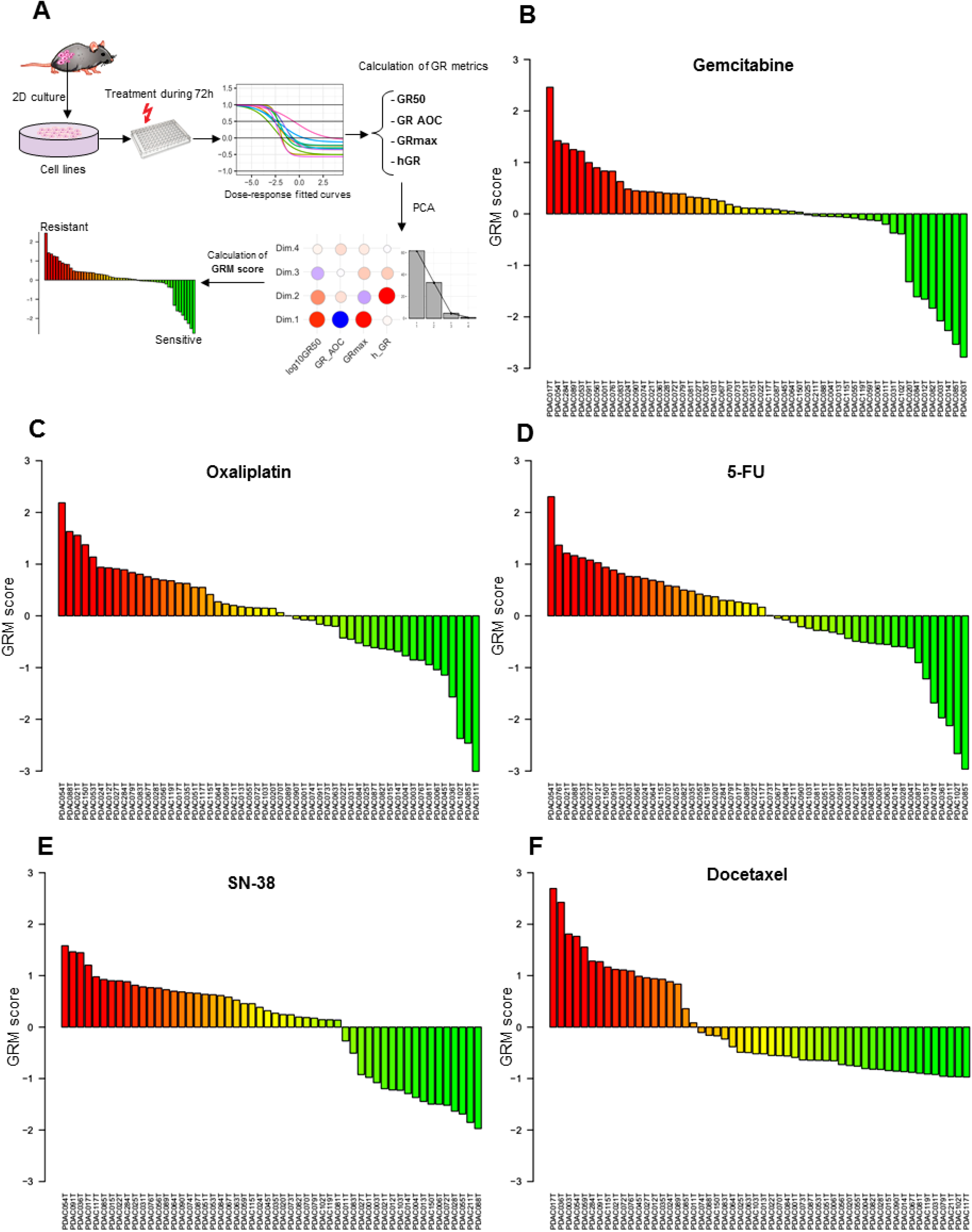
Establishment of drug resistance score. A) Schematic illustration of GRM score calculation based on Growth rate fitted curves of dose-response of PDAC cell lines treated with different drugs. B to F) Ranking of cell lines from the most resistant to the most sensitive based on GRM scores respectively for gemcitabine, oxaliplatin, 5-FU, SN-38 and docetaxel. Red to green colours reflect the gradient of resistance/sensitivity to the drug. Scores were scaled and centered around zero.

### Identification of metabolic markers associated to gemcitabine resistance

As described above, the GRM score allowed the ranking of patients according to gemcitabine resistance (from the most sensitive to the most resistant). To investigate the metabolites associated with gemcitabine resistance, we performed a *Pearson* regression analysis between all the metabolites in the dataset and gemcitabine GRM scores (correlation threshold p < 0.05). As shown in Figure 4A, the majority of identified metabolites belonged to the glycerophospholipid class, such as diacyl-PC (i.e. DAPC47 and DAPC16) and diacyl-PE (i.e. DAPE09), which positively correlated to the resistant score, whereas several monoacyl-PCs (i.e. MAPC16, MAPC07), monacyl-PE (i.e. MAPE31) and mono-phosphatidylinositols (PI), such as MAPI03, negatively correlated. In these monoacyl-PC or -PE, namely also lyso-PC or lyso-PE (LPC and LPE, respectively), one fatty acid group is removed and a phosphorylcholine or a phosphorylethanolamine moiety occupies a glycerol substitution site. In addition, amino acids, such as aspartic acid (AA13) and aminoadipic acid (AA28), were increased in resistant cells (Figure 4A and Supplementary Figure S6). However, we found a significant decrease in sterol metabolites in resistant cells, including the cholesteryl esters (ChoE_13, ChoE_14 and ChoE_17). Finally, sphingolipids, such as ceramids (Cer_02 and Cer_13) and sphingomyelins (SphLip_04, SphLip_04 and SphLip_13), also inversely correlated to gemcitabine resistance (Figure 4A and Supplementary Figure S6). These results indicate that the resistance of tumors to chemotherapy treatment with gemcitabine is associated to significant changes in metabolic profiles and alteration of several metabolites, especially lipid ones.

**Figure 4.**
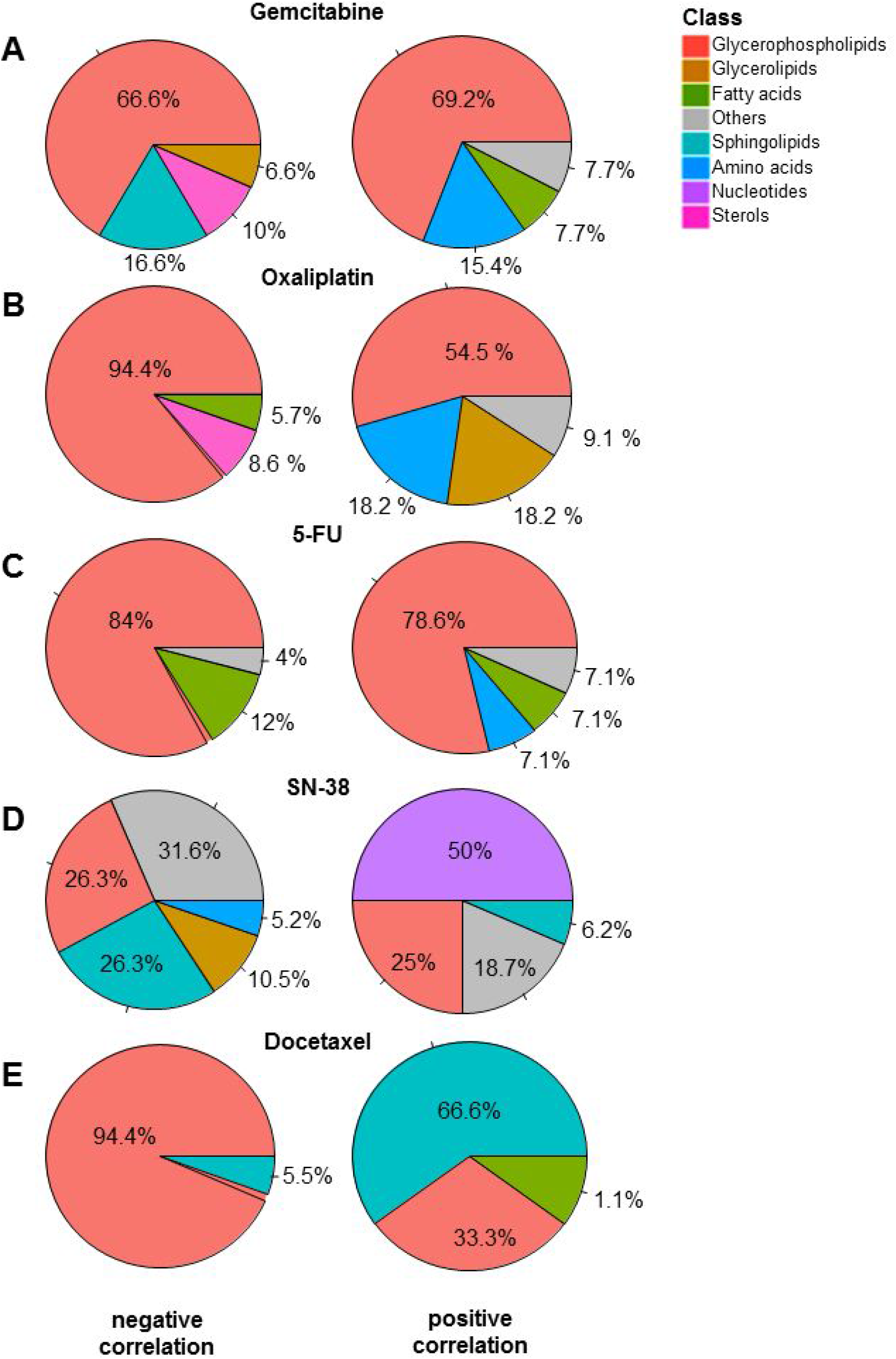
Metabolic features associated to cytotoxic drugs. Pie charts representing the negative (left) and positive (right) correlated metabolites to the score of resistance to Gemcitabine (A) Oxaliplatin (B), 5-FU (C), SN-38 (D) and Docetaxel (E).

### Metabolic features associated to oxaliplatin resistance

Similar to gemcitabine, we identified metabolites significantly correlated with oxaliplatin GRM scores. As shown in Figure 4B, most of these metabolic features associated to oxaliplatin resistance were glycerophospholipids. LPC, such as MAPC49 and MAPC45, were particularly low in resistant cells (highly anti-correlated), as opposed to DAPCs (diacyl-PC), including DAPC47 and DAPC09, along with TAGs, such as TG22 and TG26, which were positively associated to oxaliplatin resistance (Figure 4B and Supplementary Figure S7). Interestingly, similar metabolic patterns were identified for oxaliplatin and gemcitabine resistance. In fact, sterol levels (i.e. ChoE_03, ChoE_07) were low in resistant cells in conjunction with an increase of amino acids, such as aspartic acid (AA13) and hypotaurine (AA26). Beyond the singular metabolic features associated to drug resistance presented here, our results suggest that the metabolic profile that characterizes drug-resistant cells in PDAC could be similar for both oxaliplatin and gemcitabine.

### Metabolic features associated to 5-fluorouracil resistance

For analysis between metabolic features and 5-FU drug resistance, we found a negative correlation with principally LPC, such MAPC19 and MAPC38, along with some LPI, including MAPI11 and MAPI09 (Figure 4C). In addition, three free fatty acids (FFAs) decreased in resistant cells, including margaric acid (saturated FFA34), eicosenoic acid (monounsaturated FFA38) and oxidized linoleic acid (polyunsaturated FFAox35). In contrast, our results showed an increased level of a polyunsaturated fatty acid (FFA29; docosapentaenoic acid) in resistant cells to 5-FU treatment, along with hypotaurine (amino acid AA26) and fatty ester hexanoylcarnitine (AC01). Moreover, many diacyl-PC (i.e. DAPC09, DAPC46) and LPE (MAPE34, MAPE38) significantly associated with an increased resistance score to 5-FU (Figure 4C and Supplementary Figure S8). These findings again demonstrate the important association of metabolic changes and the resistance to anticancer drugs, such as 5FU, confirming the potential of standard chemotherapeutics combined with targeting cancer metabolism as promising treatment strategies for PDAC.

### Metabolic features associated to SN-38 resistance

Correlation analysis was also utilized to identify numerous metabolites that are associated with GRM scores of SN-38, an active analog of irinotecan. Particularly, we found several nucleotides that were highly associated to SN-38 resistance, including uridine 5-monophosphate (a pyrimidine nucleotide Nucl_14), ADP-glucose (purine nucleotides Nucl_22 and Nucl_23), inosine (Nucl_16), guanosine (Nucl_02) and cyclic AMP (Nucl_25) and cytidine monophosphate (Nucl_07), as shown in Figure 4E. In addition to nucleotides, some LPI, including MAPI08 and MAPI13, and MAPI05, along with a single sphingomyelin (SphLip_25) were positively associated with SN-38 resistance score. In contrast, several sphingomyelins inversely correlated with SN-38 resistance scores (Figure 4E and Supplementary Figure S9). Moreover, some PLs, such as diacy-PI DAPI06, diacyl-PE DAPE25 and MEMAPE13 (1-ether, 2-acylglycerophosphocholine), along with acylcarnitines (fatty esters), including L-octanoylcarnitine (AC02), decanoylcarnitine (AC04) and dodecanoylcarnitine (AC05), also had low levels in resistant cells. Finally, we identified a single essential amino acid (AA10, Isoleucine), which negatively associated with resistant cells. This data indicates that the metabolic reprogramming related to SN38 resistance results in accumulation of nucleotides in resistant cells, possibly generated by enhanced *de Novo* synthesis.

### Metabolic features associated to docetaxel

Investigation of metabolic features associated to docetaxel resistance showed increased levels of many sphingolipids, particularly ceramides, such as Cer_05, Cer_03 and Cer_09 (Figure 4E and Supplementary Figure S10). Eicosenoic acid (FF18) and some LPC, such as MAPC03, MAPC11 and MAPC02, also positively associated with resistance. However, almost all metabolites that negatively correlated to resistance belonged to two main subclasses of phospholipids, namely diacyl-LPE, including DAPE28 and DAPE29, and diacyl-PC, such as DAPC20 and DAPC4 (Figure 4E and Supplementary Figure S10). Thus, resistance to docetaxel appears to elicit its own unique metabolic profile.

### Multi-drug resistance metabolic features

Our exploration of metabolic features associated to drug resistance identified common metabolites that were significantly correlated in a positive manner to three out of five drugs studied here. Unsupervised clustering based on correlations between GRM scores of each drug and metabolites revealed two distinct main clusters of metabolites, indicating global positive and negative correlations to multidrug resistance (Figure 5). In fact, gemcitabine, oxaliplatin and 5-FU GRM scores clustered together, highlighting a similar profile of associated metabolites (Figure 5A and 5B). Interestingly, we found that phospholipids containing two acyl-groups, such as diacyl-PC and diacy-PE, along with some TAGs could be universally considered as metabolic markers of multidrug resistant (Figure 5A). In contrast, lysophospholipids and some cholesteryl esters demonstrated a general inverse correlation with multidrug resistance across these three drugs (Figure 5B and 5C). These results indicate that PLs, which constitute the major components of the plasma membrane of the cell, could play a key role in the acquisition of multidrug resistance in PDAC tumors.

**Figure 5.**
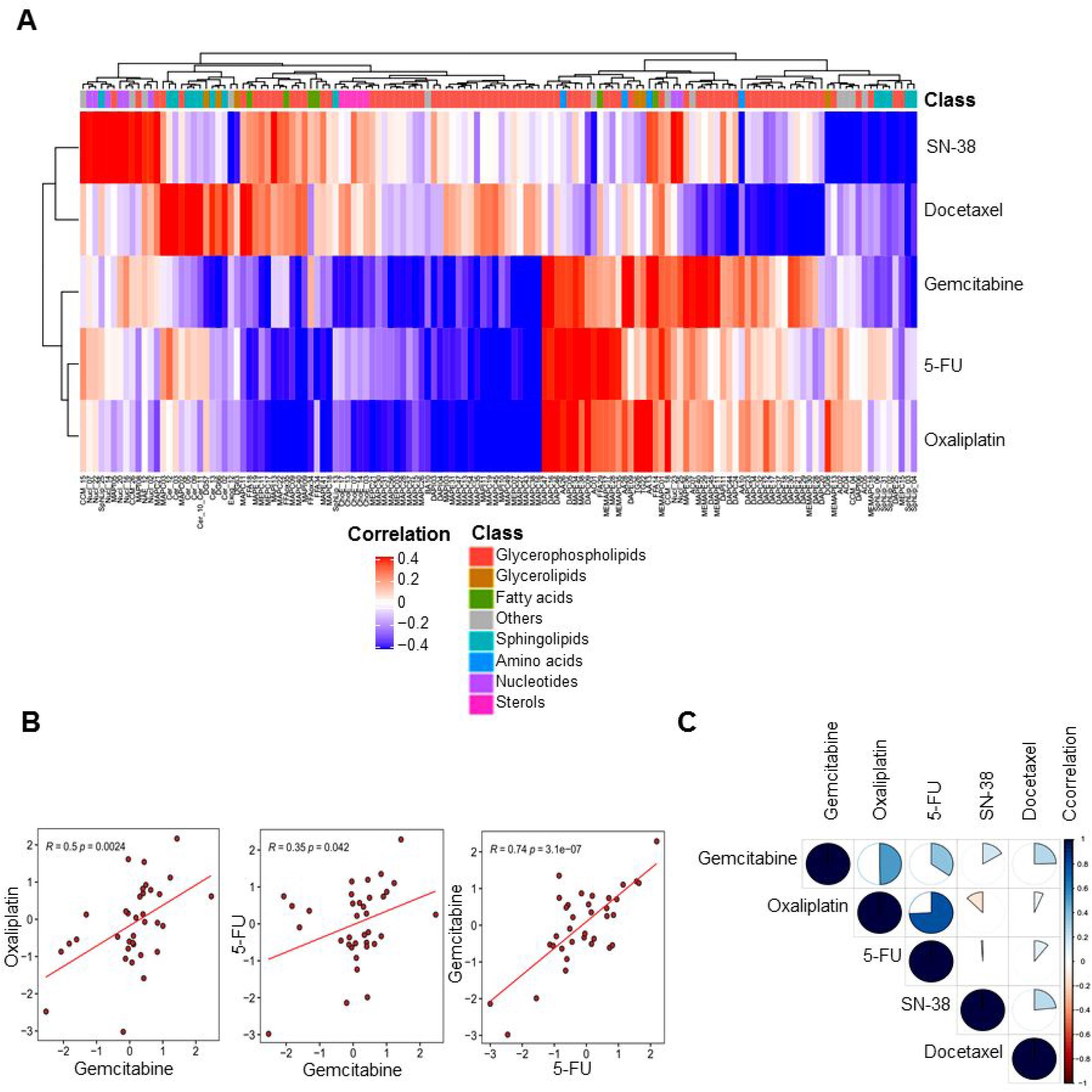
A) Heatmap of the hierarchical clustering based on the correlation of all identified metabolic features that were associated to at least one drug. Red and blue colors indicate positive and negative correlation, whereas the white color indicates the absence of correlation. B) Scatterplots comparing the GRM scores among the three significantly correlated drugs; gemcitabine, oxaliplatin and 5-FU. C) Graphic visualization of correlation matrix among GRM scores of the five drugs. The proportion of the pie chart indicates the level of correlation and the color indicates positive and negative correlation.

### Targeting the lipid profile for improving chemosensitivity to standard anticancer drugs in PDAC models

Our investigations of associations between metabolic profiles and chemoresistance offer more insights on the metabolic changes related to anticancer drug resistance. As described above, several metabolites belonging to diacyl-PC and TAGs correlated with multidrug resistance. In particular, glycerophospholipid content positively correlated to multi-drug resistance, indicating a possible role in the acquisition of this cellular phenotype. To understand whether targeting the synthesis of these metabolites could impact the response to chemotherapeutics, we treated primary PDAC-derived cells with increasing concentrations of gemcitabine, oxaliplatin or 5-FU alone or together with the specific inhibitor of glycerol 3-phosphate acyltransferase (GPAT1) FSG67. GPAT1 esterifies acyl-group from acyl-ACP to the sn-1 position of glycerol-3-phosphate, an essential step in glycerophospholipid and triacylglycerol biosynthesis. The GR_AOC in the presence or absence of FSG67 was analyzed (Figure 6). Remarkably, inhibition of glycerophospholipid synthesis is almost systematically followed by an improved sensitivity to the three cytotoxic drugs tested in several primary cultures. Although more validation experiments are required, these results indicate that the metabolism is a promising therapeutic target to overcome the challenge of chemotherapeutic resistance in pancreatic cancer cells.

**Figure 6.**
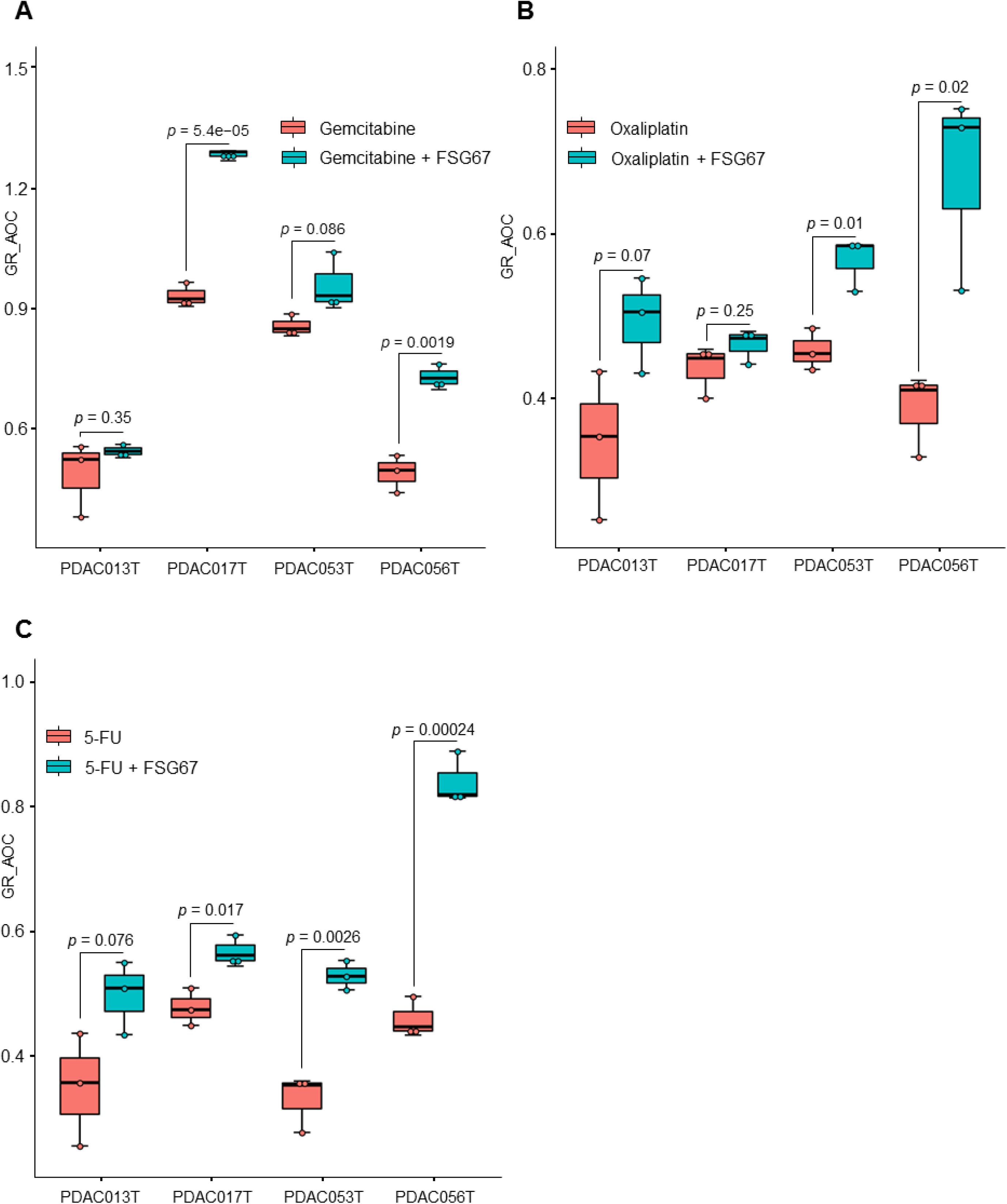
Effect of FSG67 on treatment with gemcitabine, oxaliplatin and 5-FU. PDAC013T, PDAC017T, PDAC053T and PDAC056T primary cells were treated with increasing concentrations of gemcitabine (A), oxaliplatin (B) and 5-FU (C) in the presence (green) or absence (red) of FSG67. The GR_AOC, represented as boxplots, in the presence or absence of the FSG67 was analyzed.

## Discussion

Although several investigations provide an increasing knowledge on the characterization and treatment of PDAC, it remains one of the most lethal diseases with poor prognosis and distinctive chemoresistance development. Many studies have allowed better understanding of the complexity of PDAC tumors, based mainly on molecular characterization and classification. PDAC cells demonstrate a significantly reprogrammed metabolism, which facilitates their adaptation to ensure energy and biomass sources for survival and proliferation (Guillaumond et al, 2013; Vander Heiden et al, 2009). Thus, in this study, we establish another facet to characterize PDAC tumors at the metabolic level, as a promising area of investigation to offer new insights into the association between PDAC metabolic profiles and their transcriptomic profiles, as well as their resistance to standard chemotherapeutic drugs. Our metabolic profiling of 77 PDTX samples showed that most metabolites were lipids, especially glycerophospholipids and glycerolipids. Unsupervised clustering based on metabolic profiles revealed an important heterogeneity among patients, indicating significant changes in metabolite levels. To characterize this metabolic heterogeneity among tumors, we identified a metabolic signature using ICA analysis. This signature allowed us to, first, draw significant distinctions between patients with worse versus improved clinical outcomes (Figures 1D and 1E). This finding indicates that changes in metabolic features could play a key role in tumor aggressiveness along with patient survival. In addition to its patient prognostic value, the metabolic signature identified here strongly correlated to the PAMG that describes PDAC tumors from the pure basal-like to the pure classical phenotype. PDAC with high ICA5 scores were more classical and characterized by increased levels of glycerophospholipids, while PDAC with low ICA5 scores had a basal-like phenotype with increased levels of triacylglycerols. This result is in concordance with recent studies showing that the accumulation of triacylglycerols correlates with a more aggressive phenotype in lung carcinoma cells (Nicolle et al, 2017; Tomin et al, 2018). Thus, increasing TAG lipolysis in basal-like tumors could be a favorable target to reduce the aggressiveness of tumor cells in which the accumulation of lipid droplets (containing mainly TAGs) constitute an important source of fatty acid and energy for cell growth and proliferation. Moreover, a previous study on PDAC cell lines found a low level of redox metabolites in the glycolytic phenotype, which corresponded with a basal-like cell subtype (Daemen et al, 2015). Similar results were obtained in our study regarding this subclass of metabolites, which were increased in classical tumors (high ICA5 and high PAMG). In fact, ICA5 scores of PDAC significantly correlated with redox related metabolites, such as CCM_26, which is Flavin adenine dinucleotide (*Pearson* coefficient = 0.37, p-value = 0.0032) and CCM_34, a nicotinamide adenine dinucleotide (*Pearson* coefficient = 0.3, p-value = 0.018), (Supplementary Figure S11). These differences in metabolite content, which strongly associate with patient survival and tumor phenotype, demonstrate the key role of metabolic alterations in PDAC cells. In summary, our findings provide additional insights on the characterization of metabolic profiles related to previously identified PDAC phenotypes and clinical outcomes of PDAC patients.

We also investigated the relationship between resistance of PDAC-derived cells to drugs and metabolic profiles. Our results demonstrated that resistance to cytotoxic drugs associates to a significant imbalance in both type and quantity of metabolic features. Globally, glycerophospholipids seem to be the most altered metabolites between resistant and sensitive tumors, in regards to treatment with the cytotoxic drugs gemcitabine, oxaliplatin and 5-FU. Interestingly, higher resistance was characterized by increased levels of glycerophospholipids, including diacyl-PC and diacyl-PE. Conversely, most of LPC inversely correlated with the resistant score. Previous studies have shown that breast and lung cancer cells are characterized by important quantitative and qualitative alterations of lipid concentrations in the plasma membrane, highlighting the ability to adapt to different environmental conditions to ensure proliferation and survival (Cifkova et al, 2015; Hilvo et al, 2011; Marien et al, 2015; Patterson et al, 2016). Moreover, lipid composition of the plasma membrane plays a critical role in the delivery of anticancer drugs to reach their intracellular targets via passive diffusion or active transport (Alves et al, 2016; Peetla et al, 2013). One of the mechanisms of drug resistance observed in our study could be related to reduced plasma membrane fluidity in resistant cells. In fact, increased amounts of glycerophospholipids, which constitute major components of the cell membrane, could limit drug diffusion into the cell. Although the mechanistic characterization of drug resistance still very complex, studies reporting the link between modification of plasma membrane composition and anticancer drug resistance are constantly growing (Bernardes & Fialho, 2018; Blanco et al, 2014; Brachtendorf et al, 2019; Comer et al, 2017; Kopecka et al, 2020). In this study, the metabolic features that positively associated to multidrug resistant (at least to gemcitabine, oxaliplatin and 5-FU) were glycerophospholipids, which are constituted by very long chains of fatty acids with the total number of carbon atoms greater than 40. However, down-regulated metabolites, such as LPC, have only one fatty acid chain with a total number of carbon atoms less than 30 (Supplementary Figure S12). Previously, it has been shown that glycerophospholipids containing very long fatty acyl chains are abundant in the extracellular vesicles derived from gefitinib resistant cancer patients (Jung et al, 2015). Here, we employed the GPAT1 inhibitor, FSG67, to modulate the lipid profile directly on PDAC-derived primary cultures as a proof-of-concept that influencing the lipid content modulates the therapeutic response to cytotoxic drugs. GPAT1 catalyzes the conversion of glycerol-3-phosphate and long chain acyl-CoA to lysophosphatidic acid (LPA), the rate limiting step in glycerophospholipid synthesis. We found that, in multidrug resistant cells, decreasing the glycerophospholipid synthesis through FSG67 sensitizes the majority of PDAC cells to the effect of three classical cytotoxic drugs (Figure 6).

In conclusion, in this study, we report that the metabolomic profiling of pancreatic PDTX models reveals key features driving clinical outcome and drug resistance. We also demonstrate that modifying the lipidomic profile by inhibiting the GPAT1 enzyme could be used to improve sensitivity to some cytotoxic drugs, offering a promising therapeutic target to overcome challenges related to drug resistance in PDAC.

## Material and Methods

### Metabolite extraction

Seventy-seven pancreatic cancer PDTX were obtained as previously described (Duconseil et al, 2015). Endogenous metabolic profiling experiments were measured using mass spectrometry coupled to ultra-performance liquid chromatography (UPLC-MS). Due to the wide concentration range of metabolites coupled to their extensive chemical diversity, metabolite extraction was accomplished by fractionating the pancreatic tissue samples into pools of species with similar physicochemical properties, using appropriate combinations of organic solvents as recently described (Simon et al, 2020). Thus, multiple UPLC-MS based platforms were used to analyze endogenous for the extraction of the metabolites (Simon et al, 2020).

### Data preprocessing and normalization

Raw data were processed using the TargetLynx application manager for MassLynx 4.1 software (Waters Corp., Milford, USA). A set of predefined retention time, mass-to-charge ratio pairs, Rt-m/z, corresponding to metabolites included in the analysis are considered. Associated extracted ion chromatograms (mass tolerance window = 0.05 Da) are then peak-detected and noise-reduced in both the LC and MS domains. A list of chromatographic peak areas is then generated for each sample injection. Normalization factors were calculated for each metabolite by dividing their intensities in each sample by the recorded intensity of an appropriate internal standard in that same sample, following the procedure described by Martinez-Arranz et al. (Martinez-Arranz et al, 2015). Following normalization, sample injection data were returned for manual inspection of the automated integration performed by the TargetLynx software.

### Establishment of drug resistance score

Primary cell cultures were derived from 35 PDTX samples with metabolomics data. These cell cultures were treated with increasing doses of five different drugs: gemcitabine, oxaliplatin, docetaxel, the active metabolite of Irinotecan SN-38 and 5-fluorouracil (5-FU). Dose-response data were used to calculate the Gowth Rate (GR) inhibition metrics using GRmetrics R package (Hafner et al, 2016) and to obtain the fitted curves for each drug. Thus, four metrics were considered to establish the GRM (growth rate multimetrics) score, the new score of resistance to drugs based on the following growth rate inhibition metrics: 1) GR50 correspond to the concentration at which the effect reaches 50% of growth rate; 2) GRmax represent the effect at the highest concentration; 3) GR_AOC, refers the area over the curve and 4) The hill coefficient of the fitted curve (h_GR), indicating how steep the curve is. Next, we performed a principal component analysis on these metrics using the FactoMineR (Le et al, 2008) package with five dimensions on four GR metrics including GR50, GRmax, GR_AOC and hGR. For each drug, a weighting coefficient was calculated for each metric based on its correlation with PCA dimensions and the corresponding percent of variance as follow: *coeff =* ∑_*i*_(*cor*(*GR metric, dim*(*i*)) × %*variance*(*i*)).

### Chemograms in presence of FSG67 inhibitors

Four primary PDAC cells (PDAC013T, PDAC017T, PDAC053T and PDAC056T) were cultured in SFDM and plated during 24 hours before starting the experiment at 5,000 cells per well in a 96-well plate as previously described (Fraunhoffer et al, 2020). Cells were incubated with FGS67 (30 µM), or media as control, 48h before to start the cytotoxic treatments. At this concentration we did not found significant cytotoxic effect when utilized as single treatment. Cells were then treated with increasing concentrations (0 to 1000 µM) of gemcitabine, oxaliplatin or 5-FU for 72h. Cell viability was measured with PrestoBlue (Thermo Fisher Scientific) reagent and quantified using the plate reader Tristar LB941 (Berthold Technologies). Each experiment was repeated at least three times. Values were normalized and expressed as the percentage of the control (vehicle), which represents the 100% of normalized fluorescence.

### Data analysis

All statistical and bioinformatics analysis were performed using different packages and custom scripts of the R programming language. Data were log2 transformed and centered prior to downstream analysis. Hierarchical clustering analysis was performed using ComplexHeatmap library. Independent component analysis (ICA) was performed using JADE package based on five components. Cox proportional hazard regression model from the “*survival*” package was used to perform the univariate survival analysis and determine relative risk of death associated to different factors, with a confidence interval of CI = 95%. Each metabolic feature was associated to the PAMG (Nicolle et al, 2020) and drug resistance score using a significant *Pearson’s* coefficient of correlation with a p value of correlation test < 0.05.

## Acknowledgements

We thank Fabienne Guillaumond for critically reading the manuscript. This work is part of the national program Cartes d’Identité des Tumeurs (CIT) funded and developed by the Ligue Nationale Contre le Cancer. This work was supported by INCa (Grants number 2018-078 and 2018-079), Canceropole PACA, Amidex Foundation and INSERM.

## Supplementary Figure legends

**Supplementary Figure S1.**
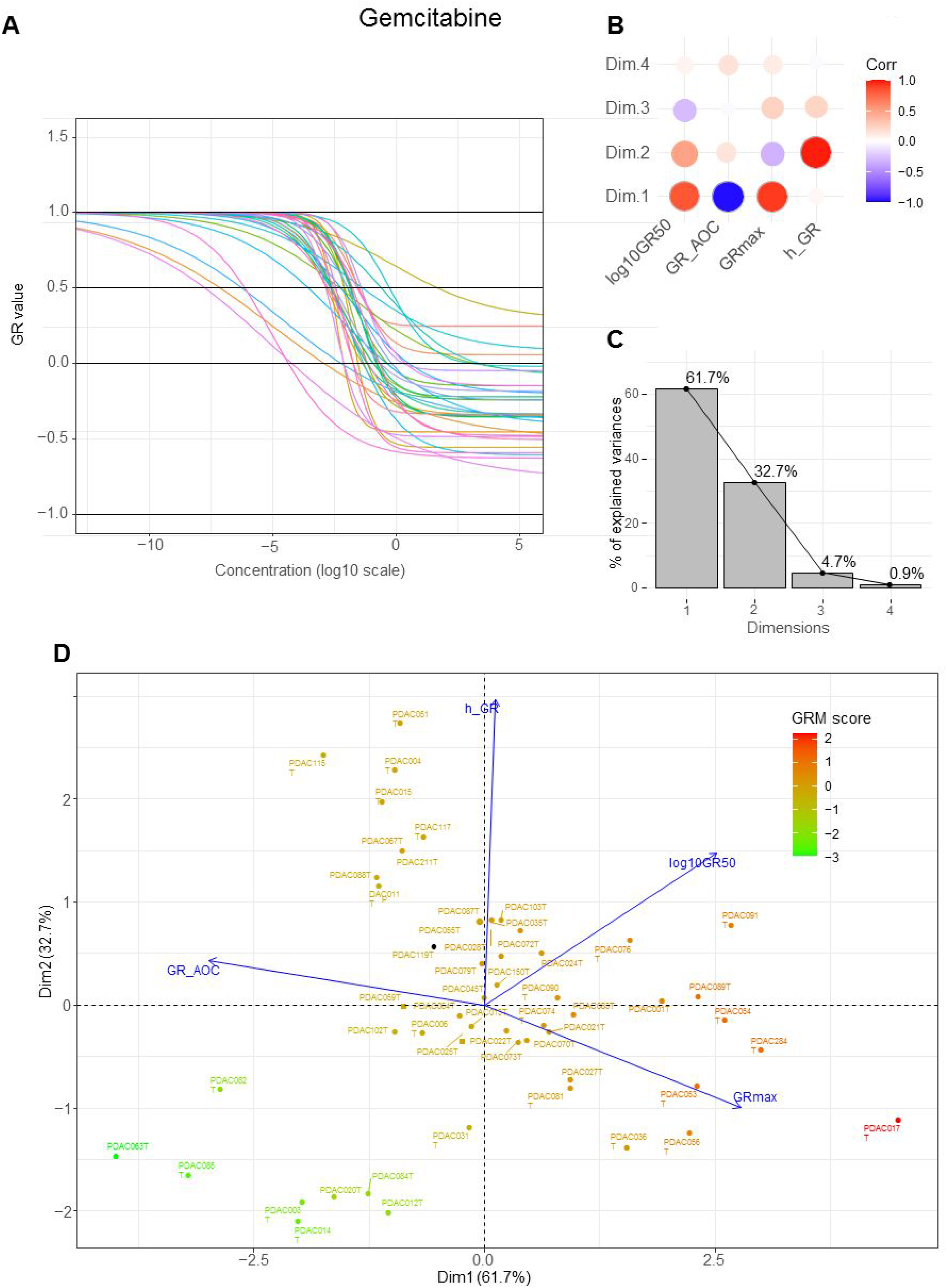
A) Fitted dose response curves of PDAC-derived primary cells treated with increasing concentrations of gemcitabine. B) Correlation of GR metrics with four principle components of the PCA. C) Barplots of the explained variance percentages by the PCA dimensions. D) Biplot of the PCA results highlighting the relationship between different GR metrics and cell lines. Red to green color reflects the GRM score.

**Supplementary Figure S2.**
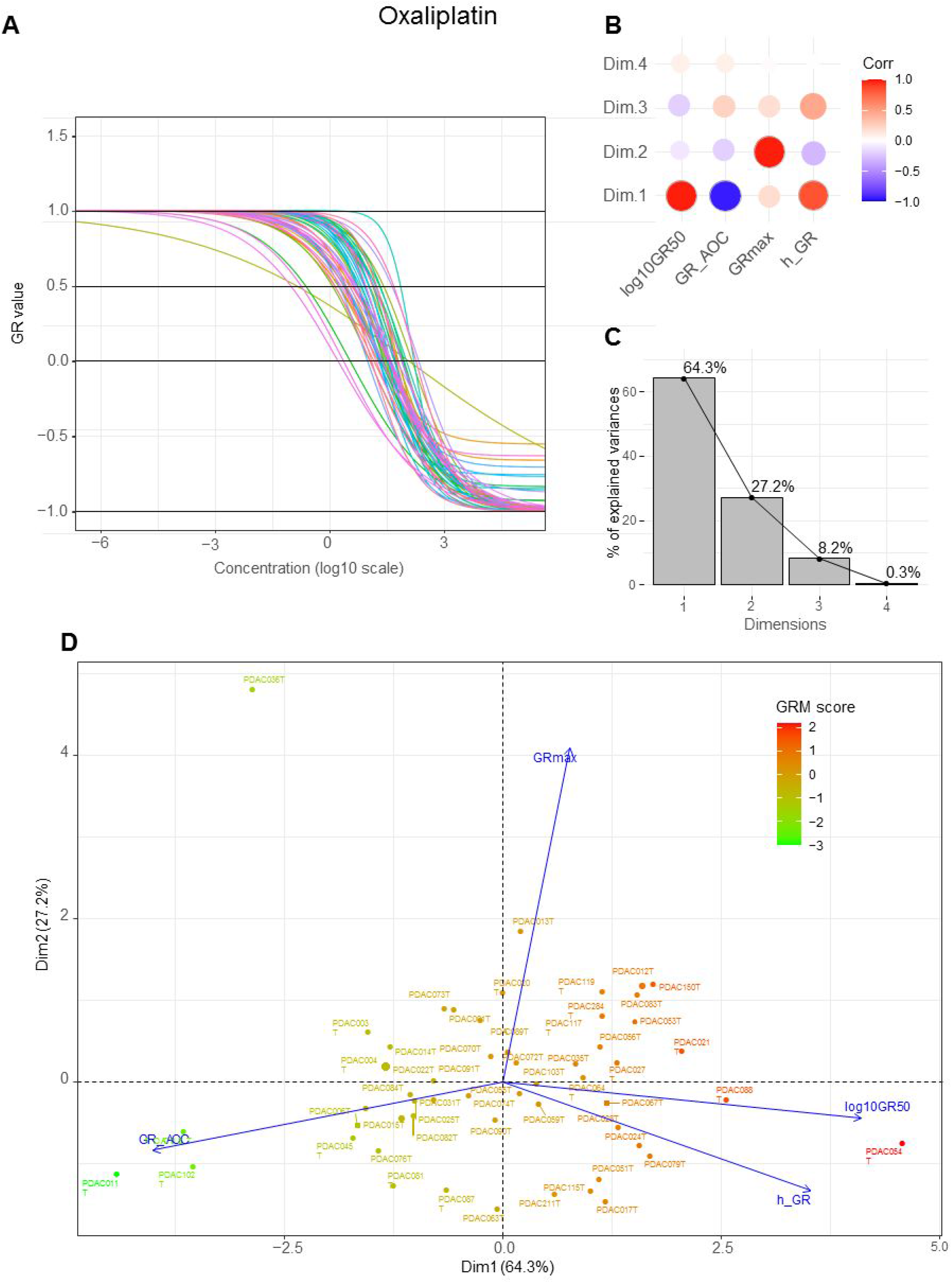
A) Fitted dose response curves of PDAC-derived primary cells treated with increasing concentrations of oxaliplatin. B) Correlation of GR metrics with four principle components of the PCA. C) Barplots of the explained variance percentages by the PCA dimensions. D) Biplot of the PCA results highlighting the relationship between different GR metrics and cell lines. Red to green color reflects the GRM score.

**Supplementary Figure S3.**
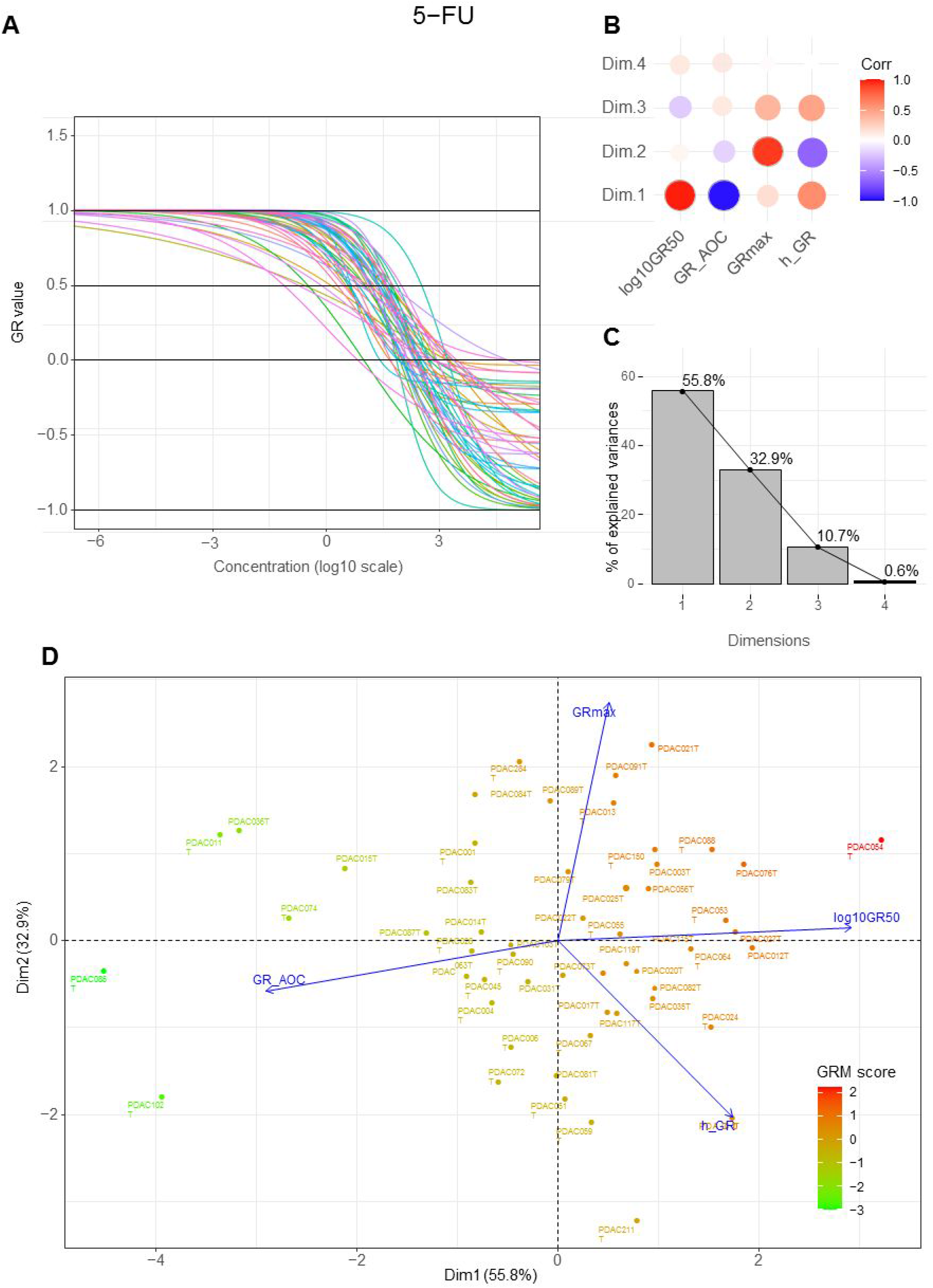
A) Fitted dose response curves of PDAC-derived primary cells treated with increasing concentrations of 5-FU. B) Correlation of GR metrics with four principle components of the PCA. C) Barplots of the explained variance percentages by the PCA dimensions. D) Biplot of the PCA results highlighting the relationship between different GR metrics and cell lines. Red to green color reflects the GRM score.

**Supplementary Figure S4.**
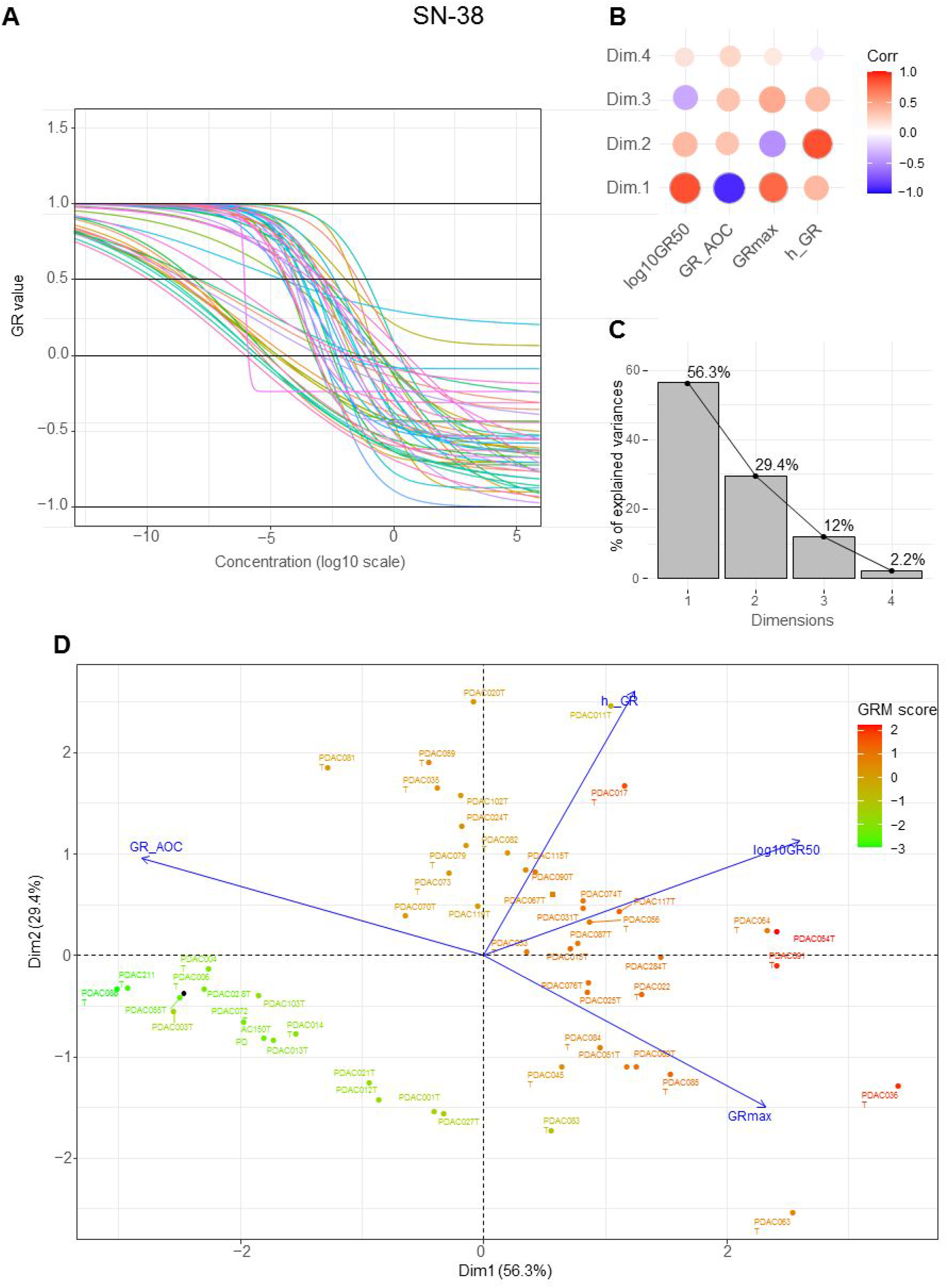
A) Fitted dose response curves of PDAC-derived primary cells treated with increasing concentrations of SN-38. B) Correlation of GR metrics with four principle components of the PCA. C) Barplots of the explained variance percentages by the PCA dimensions. D) Biplot of the PCA results highlighting the relationship between different GR metrics and cell lines. Red to green color reflects the GRM score.

**Supplementary Figure S5.**
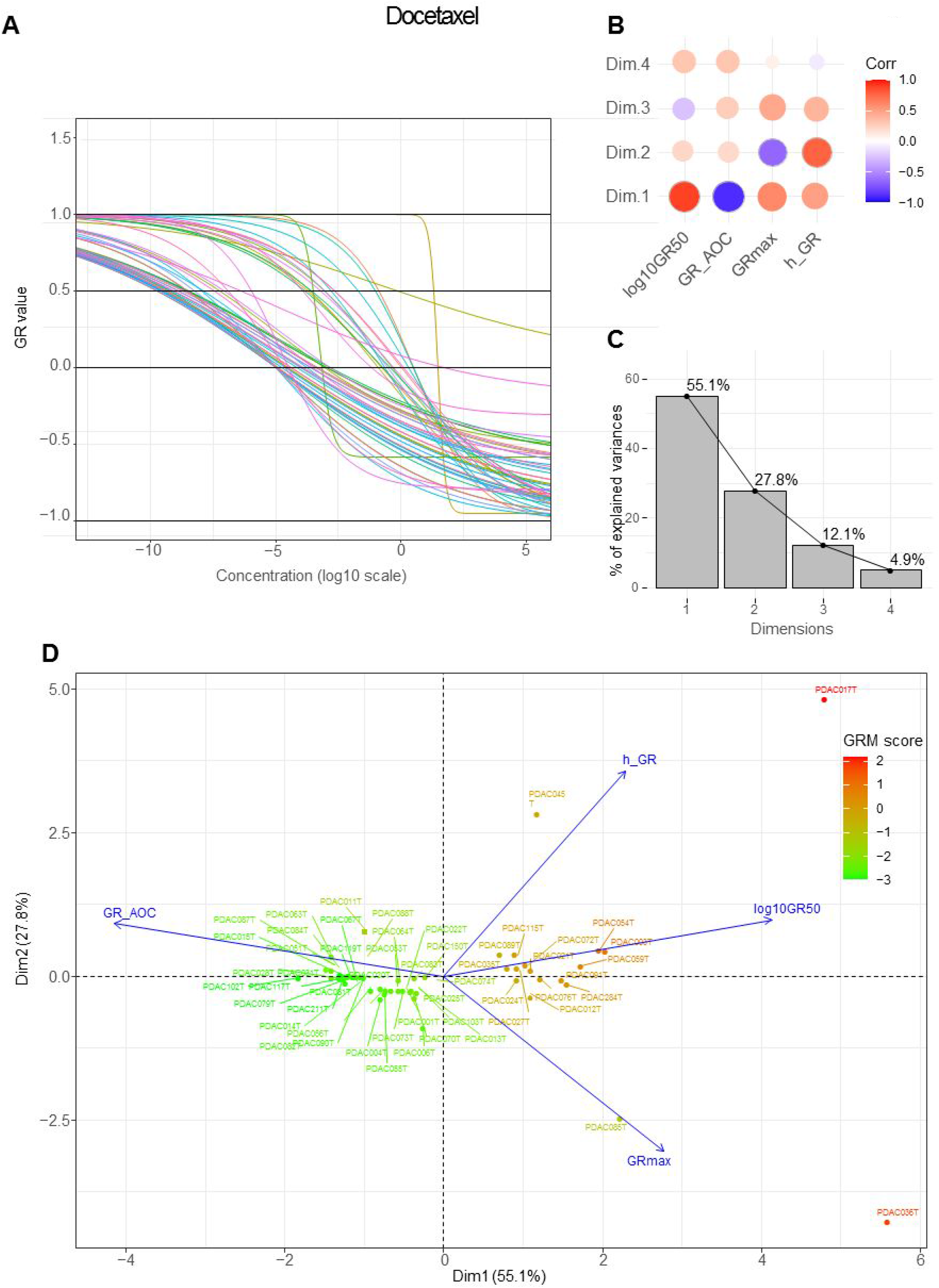
A) Fitted dose response curves of PDAC-derived primary cells treated with increasing concentrations of docetaxel. B) Correlation of GR metrics with four principle components of the PCA. C) Barplots of the explained variance percentages by the PCA dimensions. D) Biplot of the PCA results highlighting the relationship between different GR metrics and cell lines. Red to green color reflects the GRM score.

**Supplementary Figure S6.**
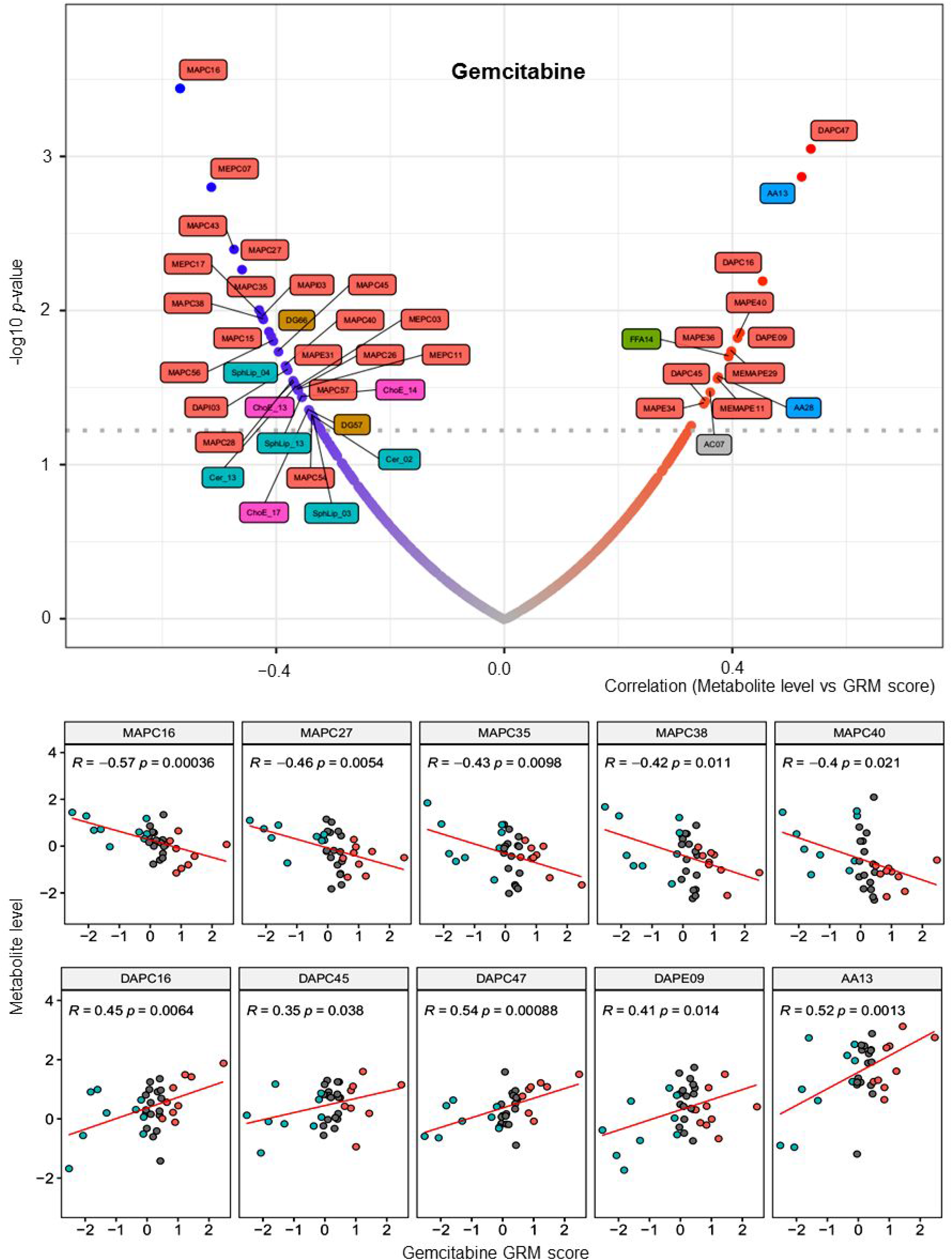
Metabolic features associated to Gemcitabine. Volcanoplot representing the negative (left) and positive (right) correlated metabolites to the score of resistance to Gemcitabine. Scatter plots of highest correlated metabolites to Gemcitabine GRM score. Statistics of the Pearson’s correlation are shown.

**Supplementary Figure S7.**
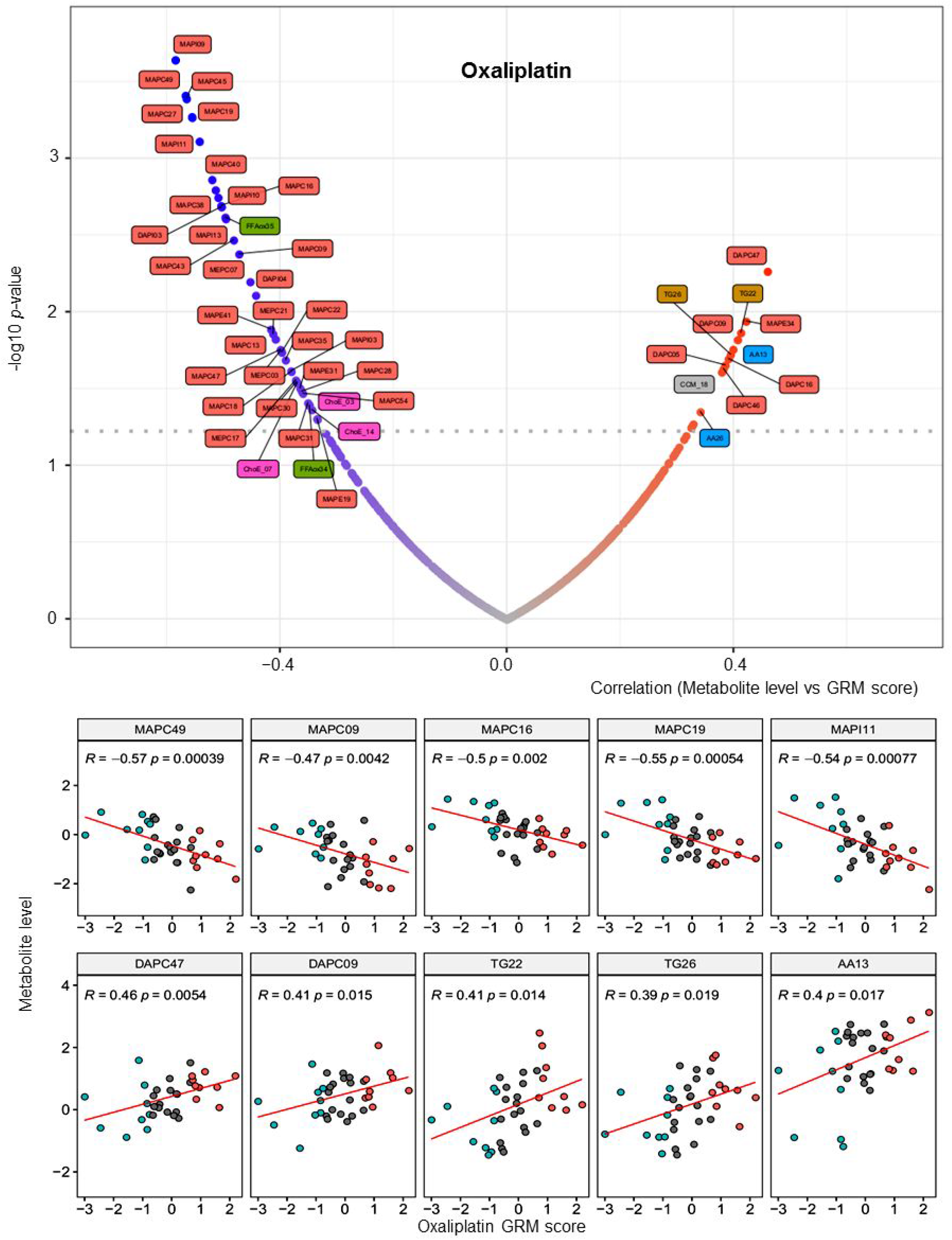
Metabolic features associated to Oxaliplatin. Volcanoplot representing the negative (left) and positive (right) correlated metabolites to the score of resistance to Oxaliplatin. Scatter plots of highest correlated metabolites to Oxaliplatin GRM score. Statistics of the Pearson’s correlation are shown.

**Supplementary Figure S8.**
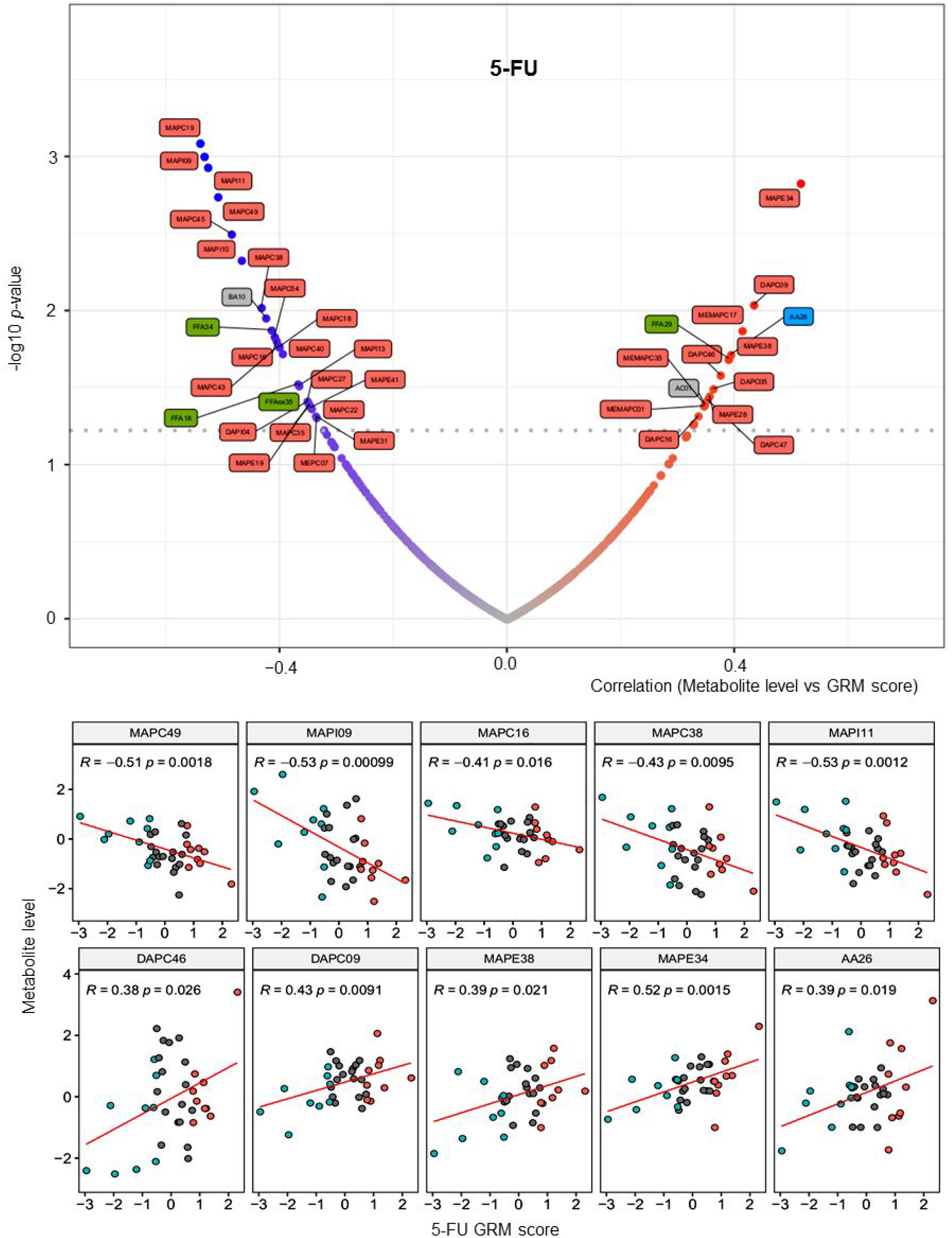
Metabolic features associated to 5-FU. Volcanoplot representing the negative (left) and positive (right) correlated metabolites to the score of resistance to 5-FU. Scatter plots of highest correlated metabolites to 5-FU GRM score. Statistics of the Pearson’s correlation are shown.

**Supplementary Figure S9.**
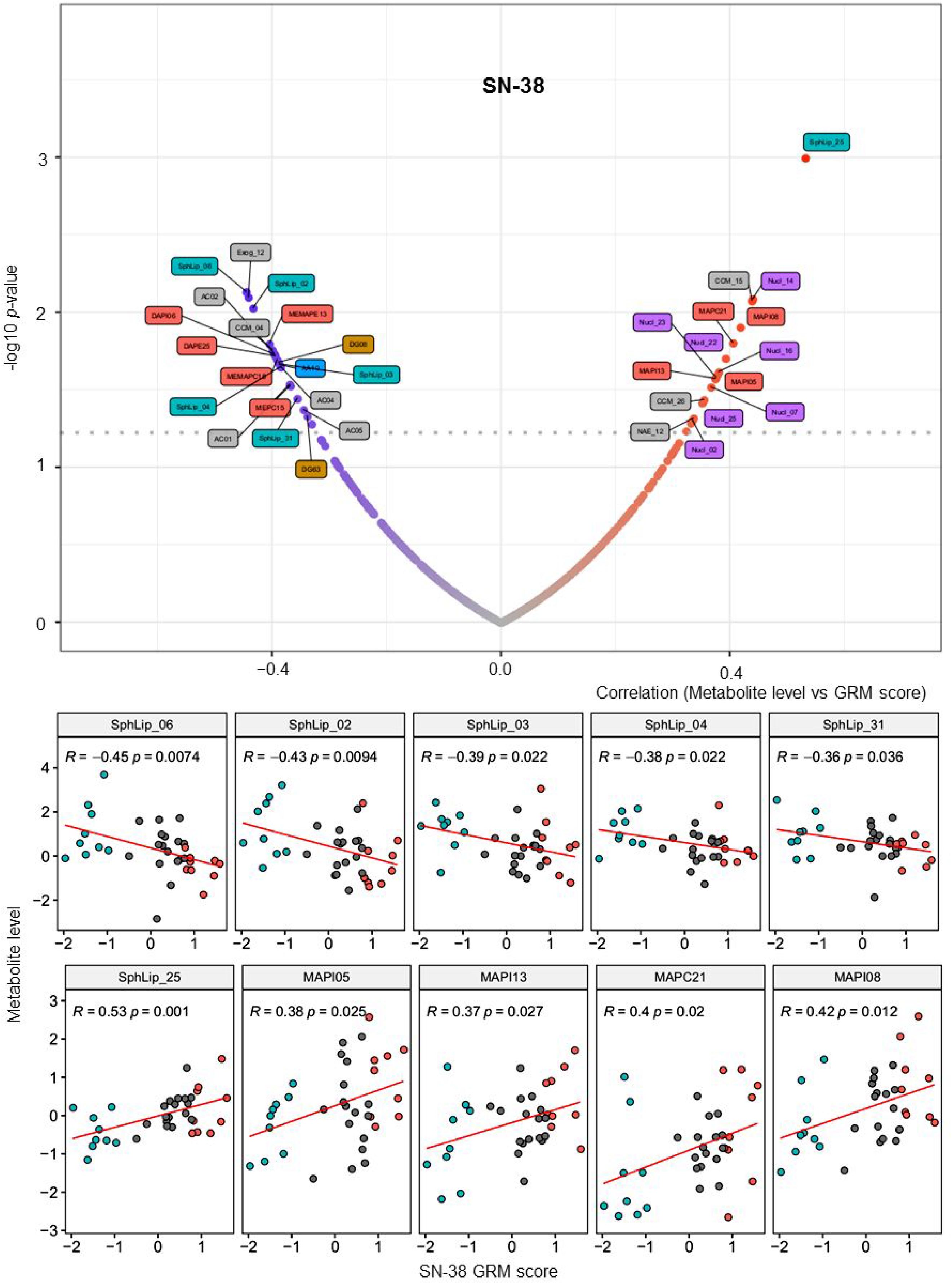
Metabolic features associated to SN-38. Volcanoplot representing the negative (left) and positive (right) correlated metabolites to the score of resistance to SN-38. Scatter plots of highest correlated metabolites to SN-38 GRM score. Statistics of the Pearson’s correlation are shown.

**Supplementary Figure S10.**
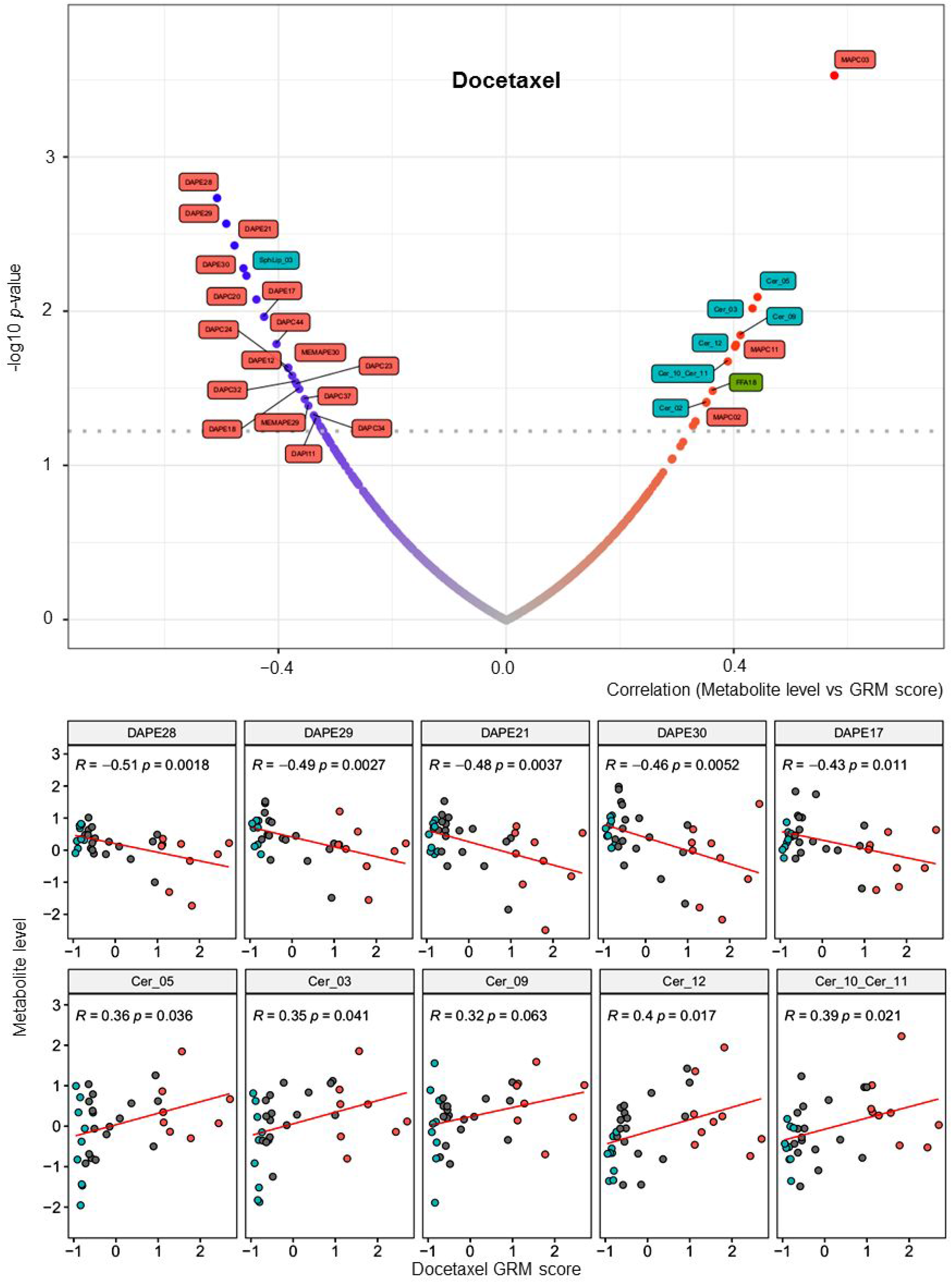
Metabolic features associated to Docetaxel. Volcanoplot representing the negative (left) and positive (right) correlated metabolites to the score of resistance to Docetaxel. Scatter plots of highest correlated metabolites to Docetaxel GRM score. Statistics of the Pearson’s correlation are shown.

**Supplementary Figure S11.**
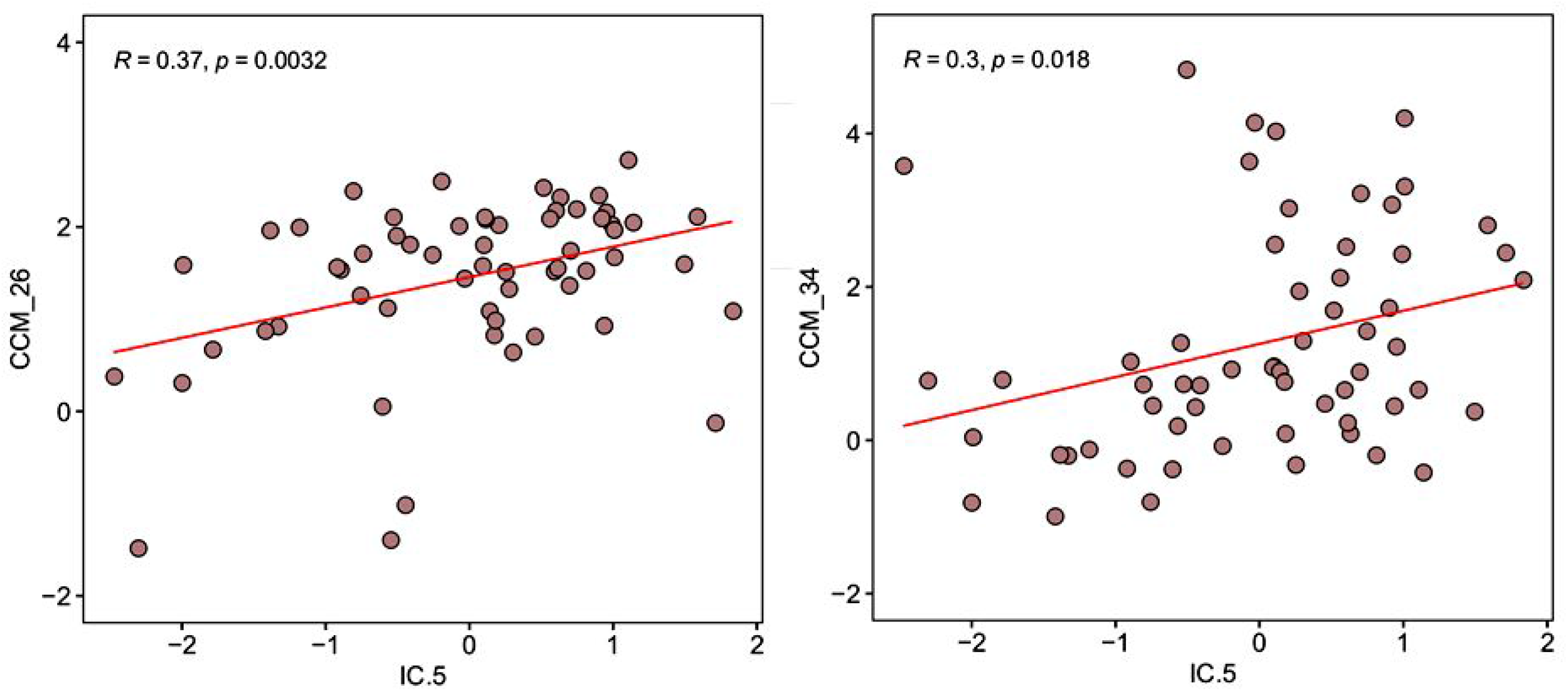
Correlation between the metabolic component ICA5 and redox associated metabolites: CCM_26 (Flavin adenine dinucleotide) and CCM_34 (nicotinamide adenine dinucleotide).

**Supplementary Figure S12.**
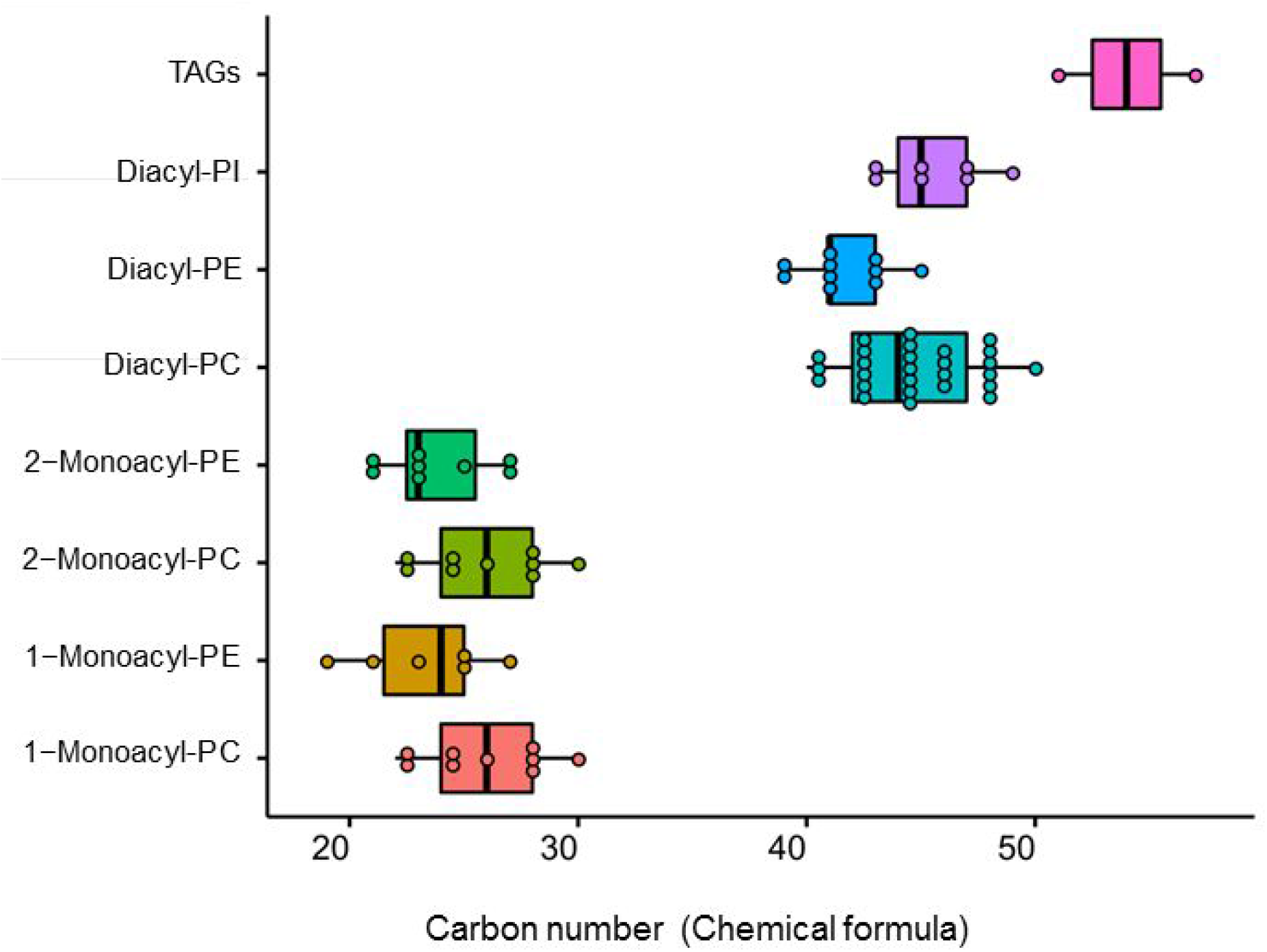
Gemcitabine, oxaliplatin and 5-FU multiresistance positively associates to glycerophospholipids that are globally constituted by very long chains of fatty acids with the number of carbon atoms greater than 40, whereas, on the contrary, those comprised of shorter fatty acid chains with the number of carbon atoms less than 30, such as lysophosphatidylcholine, were down-regulated.

